# A Rab GTPase protein FvSec4 is necessary for fumonisin B1 biosynthesis and virulence in *Fusarium verticillioides*

**DOI:** 10.1101/614206

**Authors:** Huijuan Yan, Jun Huang, Huan Zhang, Won Bo Shim

## Abstract

Rab GTPases are responsible for a variety of membrane trafficking and vesicular transportation in fungi. But the role of Rab GTPases in *Fusarium verticillioides*, one of the key corn pathogens worldwide, remains elusive. These Small GTPases in fungi, particularly those homologous to *Saccharomyces cerevisiae* Sec4, are known to be associated with protein secretion, vesicular trafficking, secondary metabolism and pathogenicity. Here, we characterized the molecular functions of FvSec4 by generating a null mutant and learned that it is important for vegetative growth, hyphal branching, and conidiation. Interestingly, the mutation did not impair the expression of key conidiation-related genes. Meanwhile, the mutant did not show any defect in sexual development, including perithecia production. GFP-FvSec4 localized to growing hyphal tips, and raised the possibility that FvSec4 is involved in protein trafficking and endocytosis. The mutant exhibited defect in corn stalk rot virulence and also significant alteration of fumonisn B1 production. The mutation led to more sensitivity to oxidative and cell wall stress agents, and defects in carbon utilization. Gene complementation fully restored the defects in the mutant demonstrating that FvSec4 plays important role in these functions. Taken together, our data indicate that FvSec4 plays important roles in *F. verticillioides* hyphal development, virulence, mycotoxin production and stresses response. Further study is needed to characterize whether the mutation in FvSec4 leads to altered vesicle trafficking and protein secretion, which ultimately impact *F. verticillioides* physiology and virulence.

## 1. Introduction

Eukaryotic cells employ exocytosis and endocytosis to ensure proper cell physiology while interacting with ambient environment (Schultzhaus and Shaw 2015). Vesicles mediate protein transport during exo- and endocytosis (Lazar, et al. 1997), and Rab GTPases play important roles in each transport step (Lazar, et al. 1997). This protein family, the largest subfamily of Ras superfamily, is involved in vesicular trafficking regulation in eukaryotes by cycling between inactive (GDP-bound) and active (GTP-bound) states (Novick 2016). In *Saccharomyces cerevisiae*, 11 members of Rab GTPases have been identified and studied (Lazar, et al. 1997). Sec4 was first identified in *S. cerevisiae* which was shown to be involved in both secretory vesicles and the plasma membrane (Salminen and Novick 1987; Goud, et al. 1988). A recent study also demonstrated that Sec4 is associated with the actin patches and endocytic internalization in yeast (Johansen, et al. 2016).

In host-pathogen interactions, pathogenic fungi such as *Fusarium* species utilize many virulence factors including cell-wall degrading enzymes, effectors and toxins by secreting these into the extracellular space or the host cytoplasm to trigger a variety of responses in the host (Ma, et al. 2013). Thus, it is reasonable to anticipate that exocyst complex plays an important role in fungal pathogenesis (Chen, et al. 2015). And Sec4 is a crucial component during this process which is responsible for the transport of post-Golgi-derived secretory vesicles to the cell membrane (Salminen and Novick 1987). In pathogenic fungi, a number of studies have shown that Sec4 is associated with various cellular functions important for virulence. *CLPT1* in *Colletotrichum lindemuthianum* was the first Sec4-like Rab GTPase gene reported in a phytopathogenic fungus associated with intracellular vesicular trafficking (Dumas, et al. 2001). Later, *CLPT1* was further demonstrated to be required for protein secretion and fungal pathogenicity (Siriputthaiwan, et al. 2005). In *Aspergillus fumigatus*, Sec4 homolog SrgA was shown to be involved in stress response, virulence and phenotypic heterogeneity (Powers-Fletcher, et al. 2013). *Magnaporthe oryzae* MoSec4 mutant exhibited defects in extracellular proteins secretion and consequently hyphal development and pathogenicity in the rice blast fungus (Zheng, et al. 2016).

*Fusarium verticillioides* (teleomorph *Gibberella moniliformis* Wineland) is a fungal pathogen of corn causing ear rot and stalk rot, posing significant food safety and security risks. The fungus is a heterothallic ascomycete, but predominantly utilizes asexual spores, *i.e.* macroconidia and microconidia, to rapidly reproduce on infected seeds and plant debris (Leslie and Summerell 2008). Most importantly, the fungus can produce various mycotoxins and biologically active metabolites including fusaric acid, fusarins, and fumonisins on infested corn ears. Fumonisin B1 (FB1) is the most prevalent and toxic form of fumonisins, a group of polyketide-derived mycotoxins structurally similar to sphinganine, and this mycotoxin is linked to devastating health risks in humans and animals, including esophageal cancer and neural tube defect (Alexander, et al. 2009; Wu, et al. 2014). Fumonisin biosynthesis gene cluster, also referred to as the *FUM* cluster, was first discovered by Proctor et al (1999). The cluster consists of a series of key genes encoding biosynthetic enzymes and regulatory proteins (Alexander, et al. 2009), and molecular characterization of *FUM1*, a polyketide synthase (PKS) gene, and *FUM8*, an aminotransferase gene, showed their important roles in FB1 biosynthesis. FB1 production was significantly reduced in *fum1* and *fum8* knockout mutants suggesting that these two key genes in the *FUM* cluster are critical for fumonisin biosynthesis (Proctor, et al. 1999; Seo, et al. 2001).

However, the regulatory mechanisms involved transport and secretion of FB1 in *F. verticillioides* remain obscure. But it is reasonable to hypothesize that key mycotoxigenic fungi employ similar mechanisms (Woloshuk and Shim 2013). A study performed in *Aspergillus parasiticus* described how vesicles, not vacuoles, are primarily associated with aflatoxin biosynthesis and export (Chanda, et al. 2009). The study also illustrated the development of mycotoxigenic vesicles under conditions conducive to mycotoxin biosynthesis. A follow-up study also demonstrated that these mycotoxigenic vesicles fuse with the cytoplasmic membrane to secrete and export aflatoxin (Chanda, et al. 2010). Exocytosis relies on a exocyst complex which has eight proteins, including Exo70p, Exo84p, Sec3p, Sec5p, Sec6p, Sec8p, Sec10p and Sec15p (TerBush, et al. 1996; Chen, et al. 2015). And Sec4 was recognized as the key component regulating the exocyst assembly (Guo, et al. 1999). We hypothesize that Sec4-like Rab GTPases in *F. verticillioides* FB1 is important for FB1 synthesis and transport. To test this, we identified a *S. cerevisiae* Sec4 homolog FvSec4 and characterized it role in *F. verticillioides* vegetative growth, virulence and FB1 biosynthesis.

## 2. Material and methods

### 2.1 Fungal strains, culture media and growth conditions

*F. verticillioides* strain 7600 was used as the wild-type strains in this study (Zhang, et al. 2018). All strains were grown and evaluated on V8 juice agar (200 ml of V8 juice, 3 g of CaCO_3_ and 20 g of agar powder per liter), potato dextrose agar (PDA, Difco) and myro agar (1g of NH_4_H_2_PO_4_, 3 g of KH_2_PO4, 2 g of MgSO4·7H_2_O, 5 g of NaCl, 40 g of sucrose and 20 g of agar powder per liter) at room temperature for 8 days. For the spore production, 5 ml sterile water was added into 8 days old V8 agar plates. Spore suspensions were harvested by passing through miracloth (EMD Millipore) and counted using the hemocytometer. Newly harvested microconidia were suspended in 0.2x potato dextrose broth (PDB) for 5h and 6.5 h with gentle shaking to assay spore germinations. For genomic DNA extraction, strains were grown in YEPD liquid medium (3 g yeast extract, 10 g peptone and 20 g dextrose per liter) at 25 °C in a rotatory shaker for 4 days. For stress assays, strains were cultured on Czapek-Dox agar (2 g/L NaNO_3_, 0.5 g/L MgSO_4_-7H_2_O, 0.5 g/L KCl, 10 mg/L 14 FeSO_4_-7H_2_O, and 1 g/L K_2_HPO_4_, PH 7) amended with 70 mg/L Congo red, 2 mmol/L H_2_O_2_ or 0.01% SDS. For carbon utilization assay, 4 μl of 1 × 10^6^ conidial suspension was inoculated on the center of Czapek-Dox agar plates with four different carbon sources, *i.e.* sucrose (30 g/L), dextrose (10 g/L), fructose (10 g/L) and sorbitol(91 g/L), and incubating 8 days at room temperature. For the mating study, conidia from wild-type, ΔFvsec4, ΔFvsec4-com strains were harvested from culture grown on V8 agar for 7 days, and subsequently spread onto *F. verticillioides* strain 7598, which was grown on carrot plates following our standard protocol (Sagaram and Shim 2007).

### 2.2 Gene deletion and complementation, polymerase chain reaction (PCR), and transformation

The constructs for *F. verticillioides* transformation were generated following our laboratory standard procedures (Sagaram and Shim 2007). Briefly, DNA fragments corresponding to 5’ and 3’ flanking regions of the gene were amplified from the wild-type genomic DNA. Meanwhile, hygromycin B phosphotransferase (*HPH*) gene in pBP15 plasmid was used to obtain the HP and PH fragments. 5’ and 3’ flanking region fragments were fused with PH and HP fragments by single-joint PCR respectively (Yu, et al. 2004; Sagaram and Shim 2007). We used protoplast preparation and transformation protocols previous described in Zhang et al (2018). We used PCR to verify targeted gene deletion mutations using primers OF/OR, UAF/YG/F, UAR/GE/R (Table S1), followed by Southern blot and qPCR for further validation.

For gene complementation, wild-type *FvSEC4* gene with its native promoter was co-transformed with a geneticin-resistant gene (*GEN*) into mutant protoplasts. All transformants were screened by PCR. All primers used in this study are presented in (Table S1). To construct the GFP-FvSec4 plasmid, *FvSEC4* coding region from *F. verticillioides* cDNA was amplified. *FvSEC4* native promoter and terminator were amplified from wild-type DNA. GFP was amplified from PKNTG plasmid (Dr. Wenhui Zheng, Fujian Agriculture and Forestry University, China). These four purified products were introduced to the *Hind*III and *BamH*I sites of PKNTG using In-Fusion-HD cloning kit (Takara Bio USA). The plasmid was then sequenced and introduced into the ΔFvsec4 strain for genetic complementation and localization study.

### 2.3 Nucleic acid manipulation and Southern blot

Bacterial plasmid DNA was isolated with Wizard miniprep DNA purification system (Promega). Fungal genomic DNA isolation and Southern blot analysis were performed following standard procedures (Sambrook 2001). Briefly, 10 μg genomic DNA of each strain was completely digested with *Cla*I and probed with a ^32^P-labelled DNA fragment amplified from *F. verticillioides* genomic DNA with primers FvAF1 and FvAR1 (Table S1).

### 2.4 Corn infection and fumonisin B1 assays

Infection assays on corn seedling were conducted as previously described with minor modifications (Kim, et al. 2018). In this study, we used silver queen hybrid seeds (Burpee) for seedling inoculation with fungal spore suspension. The seedlings were collected and analyzed after a one-week growth period in the dark room. At least three biological and three technical replicates were performed for each fungal strain.

For FB1 and ergosterol extraction, four silver queen seeds were surface sterilized using the method previously described (Christensen, et al. 2012) with a minor modification. Sodium hypochlorite (6%) was replaced with 10% bleach. Next, sterilized kernels were put on autoclaved 90-mm Whatman filter paper. A scalpel was used to create wounds on the endosperm area, and these seeds were placed in 40-ml borosilicate glass vials. Fungal spore suspensions (200 μl, 10^7^/ml) were inoculated into each vial, and these samples were incubated in room temperature for 7 days. FB1 extraction and sample purification methods were described previously (Christensen, et al. 2012). HPLC analyses of FB1 and ergosterol were performed as described (Shim and Woloshuk 1999). FB1 levels were then normalized to ergosterol contents. The experiment was repeated twice with three biological replicates.

### 2.5 RNA extraction and gene expression study

A 1ml fungal spore suspension (10^6^ spores/ml) was inoculated in 100 ml YEPD for 3 days at room temperature with agitation (150 rpm). Then, mycelium from each flask was filtered through Miracloth and weighed (0.3 g) before transferred to 100 ml myro liquid medium. Samples were collected after 7 days incubation at room temperature with agitation (150 rpm). In addition, 2 ml of spore suspension (10^6^ spores/ml) were added into 100 ml YEPD, incubated for 20 h at 28°C at 150rpm before being harvested for RNA extraction for qPCR assay. Three replicates were performed for each strain. Total RNA was extracted using Qiagen kit following the manufacturers’ protocols. RNA samples were converted into cDNA using the Verso cDNA synthesis kit (Thermo Fisher Scientific), and qPCR analyses were performed with Step One plus real-time PCR system using the DyNAmo ColorFlash SYBR Green qPCR Kit (Thermo Fisher Scientific) with 0.5 μl cDNA as the template in per 10 μl reaction. Expression levels were normalized with *F. verticillioides* β-tubulin-encoding gene (FVEG_04081).

### 2.6 Microscopy and staining protocol

For hyphal growth imaging, strains were cultivated on 0.2xPDA for three days, and a block of agar containing the growing edge (approximately 1 cm diameter) was cut and placed on a glass slide. Next, water (10 μl) was added on the agar block, and subsequently a coverslip was gently placed on top. We incubated the sample at 28°C for additional 20 mins, and then observed hyphal growth under a microscope (Olympus BX60). For GFP assay, we followed a previous method with minor modifications (Schultzhaus, et al. 2015) and with assistance from Dr. Brian Shaw (Department of Plant Pathology and Microbiology, TAMU). We used 16-18 h hypha to take images and water was added on the top of agar. Samples were incubated at 28°C for 20 mins. For FM4-64 staining, 16-18h growth of strains in 0.2xPDA were cut and put in the slide. Then, we added 10 μl of 5 μM FM4-64 on top of medium and incubated at room temperature for 10 mins. We used 0.2xPDB broth to wash samples twice before taking images. Images were prepared using ImageJ software (Schneider, et al. 2012).

## 3. Results

### 3.1 Identification of the Sec4 homolog in *F*. *verticillioides*

We used the Sec4 protein sequence from the *S. cerevisiae* genome database (http://www.yeastgenome.org/) to conduct a search into NCBI *F. verticillioides* database. This search identified FVEG_06175 locus, a 990-bp gene encoding a putative 203-amino-acid protein, which was designated as *FvSEC4* gene. To identify Sec4 homolog in other fungal species, the predicted FvSec4 amino acid sequence from the *F. verticillioides* genome database was used for our BLAST search. Multiple sequences alignment (Fig. 1) and phylogenetic analyses (Fig. S1A) indicated that Sec4-like proteins share high amino acid identity in fungi, such as *S. cerevisiae* Sec4 (YFL005W, 64 % identity), *A. fumigatus* AfSrgA (Afu4g04810, 87% identity), *M. oryzae* MoSec4 (MGG_06135, 88% identity), *C. lindemuthianum* CLPT1 (AJ272025, 95% identity) and *C. orbiculare* CoSec4 (Cob_13201, 95% identity). The predicted ScSec4 protein structure was obtained from PDB (PDB ID: 1G16) and modified into alignment by ESPript (Stroupe and Brunger 2000; Robert and Gouet 2014).

**Fig. 1.**
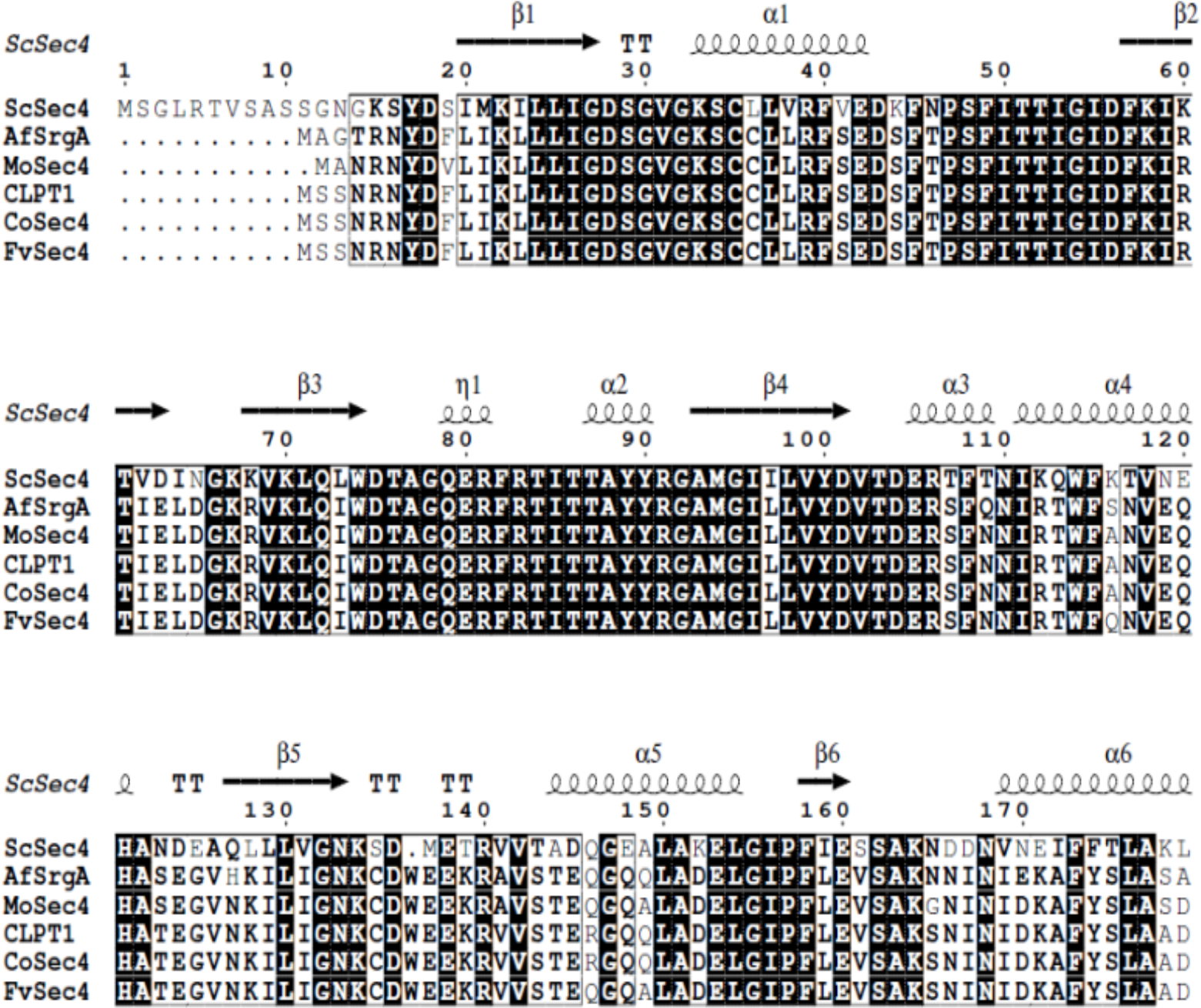
FvSec4 protein sharing high similarity with other fungal species. Protein sequence alignment of *S. cerevisiae* Sec4, *F. verticillioides* FvSec4, *A. fumigatus* AfSrgA, *M. oryzae* MoSec4. *C. lindemuthianum* CLPT1. White characters with black background and black characters in a box indicate identical and similar sequences, respectively. Sec4 is predicted to have six alpha helices and six beta strands.

### 3.2 Loss of FvSec4 impairs hyphal growth and conidiation

To investigate the function of FvSec4, we generated deletion mutants by replacing the entire gene with a hygromycin-resistance marker (Fig. S2A). The gene-replacement mutants were confirmed by Southern blot (Fig. S2B). The wild-type strain showed a 2.4-kb hybridizing band and all three putative mutants had a 6.5-kb band, as expected when using the ORF 5’ flanking probe for Southern blot (Fig. S2B). These results suggested that the three mutant strains had a single-copy insertion of the hygromycin-resistance marker and had no ectopic insertion events. We selected the first mutant, which was designated ΔFvsec4, to perform further experiments, including qPCR and phenotypic analyses in this study (Fig. S2C). We also generated a gene complementation strain ΔFvsec4-Com and a GFP-tagged complementation strain ΔFvsec4-GFP-FvSec4.

To study the role of FvSec4 protein in *F. verticillioides* vegetative growth, we cultivated wild-type, ΔFvsec4, ΔFvsec4-Com and ΔFvsec4-GFP-FvSec4 strains on V8, 0.2xPDA, myro agar media. The ΔFvsec4 mutant showed a drastic reduction in growth and less fluffy mycelia on all agar media tested. Both ΔFvsec4-Com and ΔFvsec4-GFP-FvSec4 strains exhibited full recovery of growth defects (Fig. 2A). Moreover, ΔFvsec4 displayed hyphal hyperbranching under microscopic examination (Fig. 2B). In addition, ΔFvsec4 mutant produced significantly lower quantity of conidia when compared with wild-type and complemented strains (Fig. 3A). However, conidia germination rate in the mutant and wild-type did not show a significant difference when all strains were cultivated in 0.2xPDB liquid culture (Fig. 3B). Also, mycelial fresh weight did not differ between wild-type and mutants after growing in YEPD liquid medium for 3 days (Fig. 3C).

**Fig. 2.**
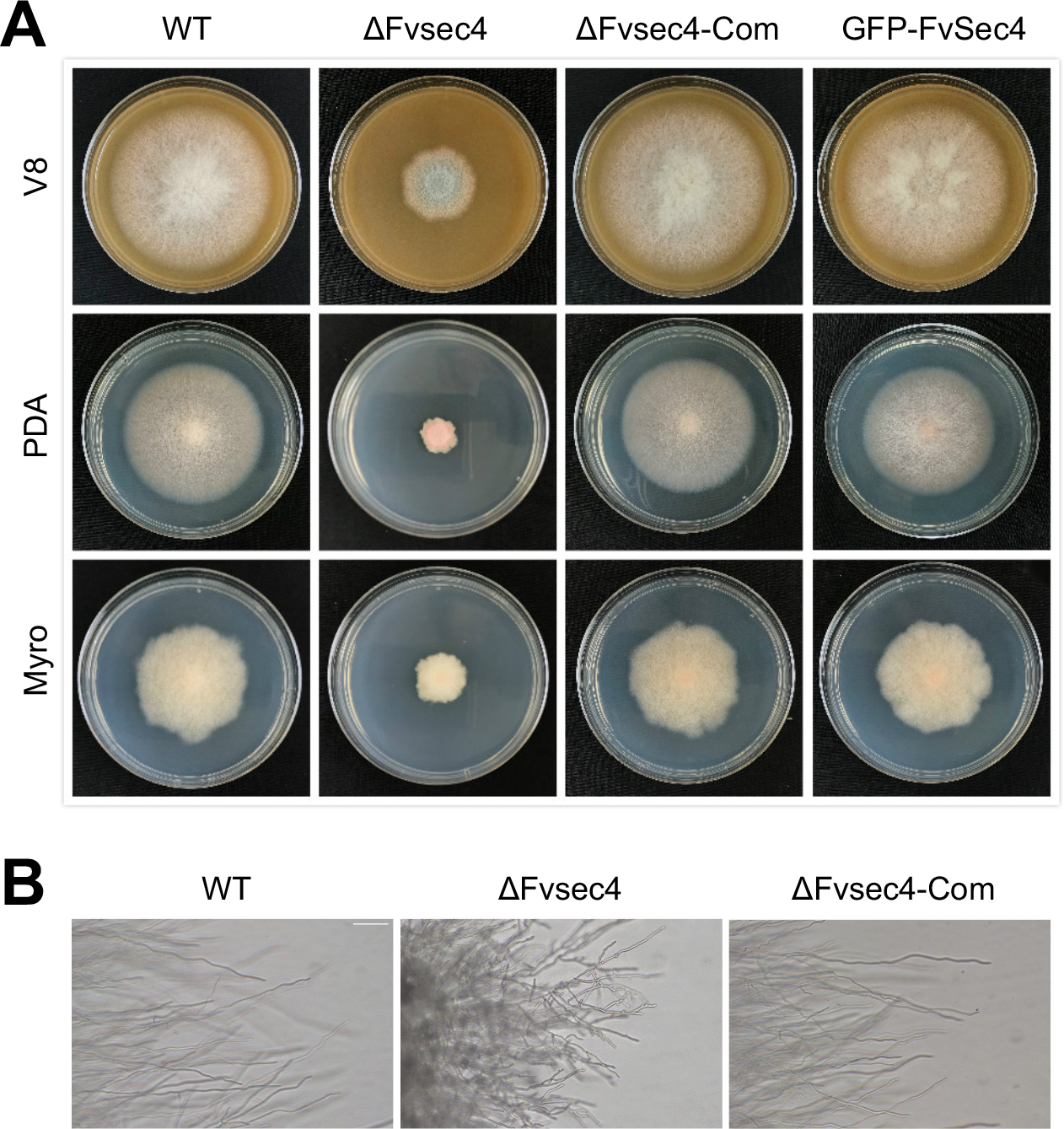
Vegetative growth of wild-type (WT), ΔFvsec4, ΔFvsec4-Com and ΔFvsec4-GFP-FvSec4 (GFP) strains. (A) Strains were cultured on the V8, 0.2xPDA, myro agar plates for 8 days at room temperature. (B) ΔFvsec4 mutant enhanced hyphal branching compared to WT, ΔFvsec4-Com after 3 days on 0.2xPDA agar plates. Bar = 100 μm

**Fig. 3.**
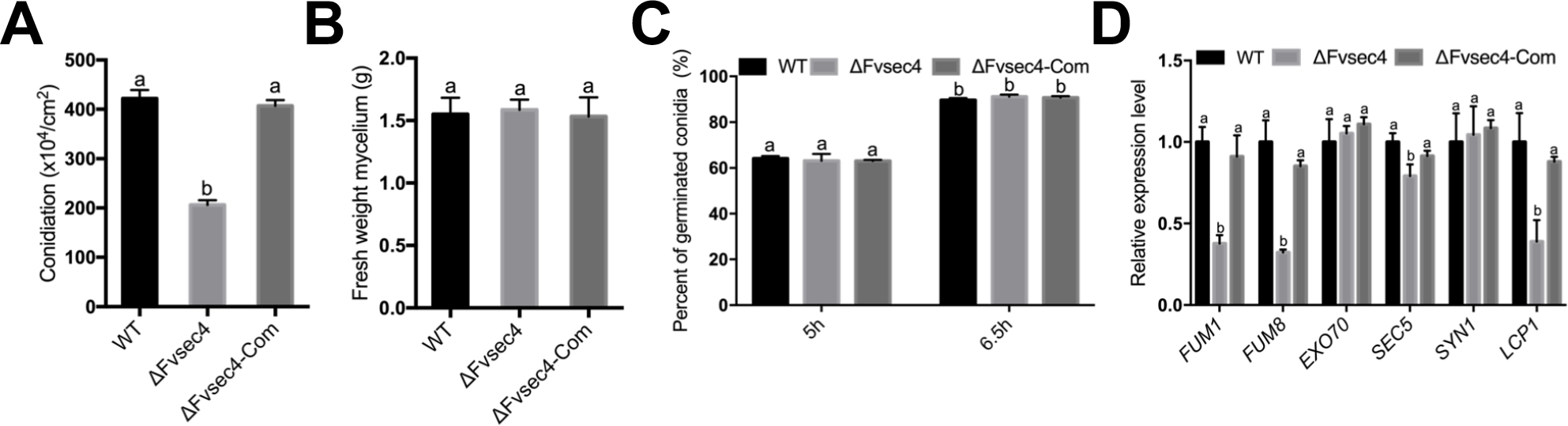
Impacts of FvSec4 on conidia production (A) Conidia production in wild-type (WT), ΔFvsec4, ΔFvsec4-Com strains were measured after incubation on V8 agar medium at room temperature for 8 days. (B) Conidia germination rate in WT, ΔFvsec4, and ΔFvsec4-Com on 0.2xPDB were examined under microscope after 5 and 6.5h incubation with gently shaking. (C) Mycelium fresh weight were assayed after 3-day incubation in YEPD liquid medium. (D) Transcript differences of conidia-related genes in ΔFvsec4 were compared to WT and ΔFvsec4-Com strains. Three biological replicates were performed independently. Error bars in this study all represent the standard deviations for three replicates. Lowercase letters in this study on the bar top suggest significant differences among various strains. (Student T-test, P<0.05). All data in this study were analyzed by Prism software.

To further characterize the basis for conidia production deficiency in ΔFvsec4, we used qPCR to test transcription levels of key conidia regulation genes, including *BRLA*, *WETA*, *ABAA* and *STUA.* Total RNA samples were extracted from strains cultured in myro broth for 7 days and in YEPD broth for 20 h. The qPCR data suggested that conidia regulation genes are not impacted by the FvSec4 deletion both in myro and YEPD culture, except *ABAA* expression that showed 40% reduction when the mutant was cultured in myro broth (Fig. S3C and 3D). We also tested whether FvSec4 is important for sexual reproduction, but all strains showed no defect perithecia production (Fig. S3E).

### 3.3 Subcellular localization of FvSec4 suggests its role in vesicle trafficking

Since Sec4 is known as a key component that regulates the exocyst assembly, we studied the localization of FvSec4 in *F. verticillioides*. We used the native promoter and fused GFP to the N-terminus of *FvSEC4* coding sequence, and subsequently transformed the *FvSec4pro*-*GFP*-*FvSec4* construct into ΔFvsec4 strain. After confirming construct insertion through PCR, we performed a live-cell imaging study of GFP-FvSec4 protein subcellular localization. The FvSec4 green fluorescent signal accumulated mainly in the tips of hyphal, which was consistent with the predicted Sec4 protein role in the tip vesicle transport and growth (Fig. 4A).

**Fig. 4.**
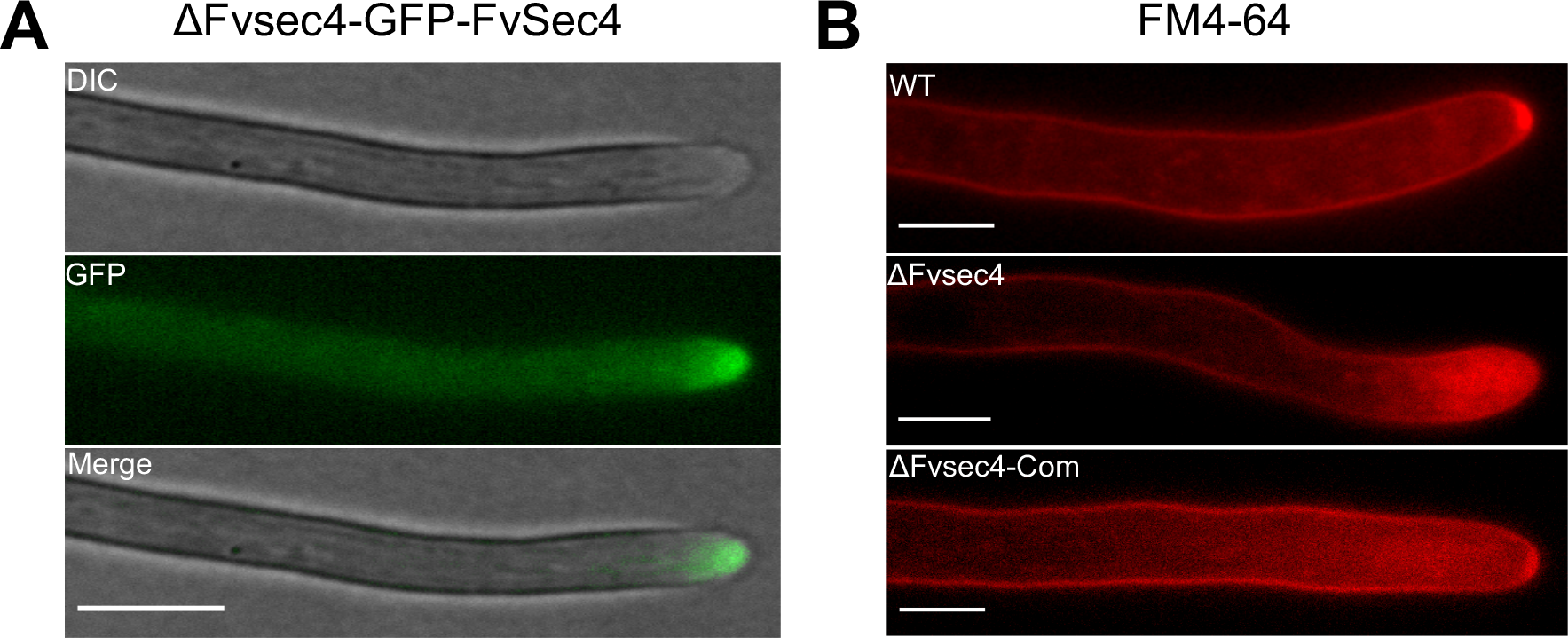
FvSec4 protein localization assay. (A) GFP-Sec4 protein driven by its native promoter mainly localized to the apical area of growing hyphae. Bar = 5μm (B) Hyphal growth of wild-type (WT), ΔFvsec4, ΔFvsec4-Com were examined under a microscope after 10 min of FM4-64 staining. The Spitzenkörper region in ΔFvsec4 was compared with WT and ΔFvsec4-Com. Bar = 5μm

In addition to exocytosis, a previous study revealed that Sec4 is also important for endocytosis (Kean, et al. 1993). To further test whether FvSec4 is associated with endocytosis, we stained the mycelia of wild-type, ΔFvsec4, and ΔFvsec4-Com with FM4-64, which is frequently used to study endocytosis and vesicle trafficking in the fungal hyphae (Fischer-Parton, et al. 2000). Strong fluorescent signals were detected in the Spitzenkörper region in both WT and ΔFvsec4-Com, while the ΔFvsec4 mutant showed a broader diffused staining at the mycelial tip. No Spitzenkörper structure staining was observed in ΔFvsec4. This result suggests that ΔFvsec4 is either not properly functioning in the uptake of FM4-64 or defective in recycling dyes to hyphal tip by exocytosis (Fig. 4B).

### 3.4 FvSec4 is important for corn seedling rot virulence

To test whether FvSec4 plays a role in *F. verticillioides* virulence, we inoculated wild-type, ΔFvsec4, ΔFvsec4-Com spore suspensions, with sterilized distilled water as the negative control, on one-week-old corn (silver queen hybrid) seedlings. After one week of incubation, we observed significantly reduced rot symptoms in ΔFvsec4 mutant when compared to the wild-type strain (Fig. 5A and 5B). Gene complementation strain ΔFvsec4-Com showed fully recovered stalk rot symptoms in our assay. These results demonstrated that FvSec4 plays an important role in *F. verticillioides* corn stalk rot virulence.

**Fig. 5.**
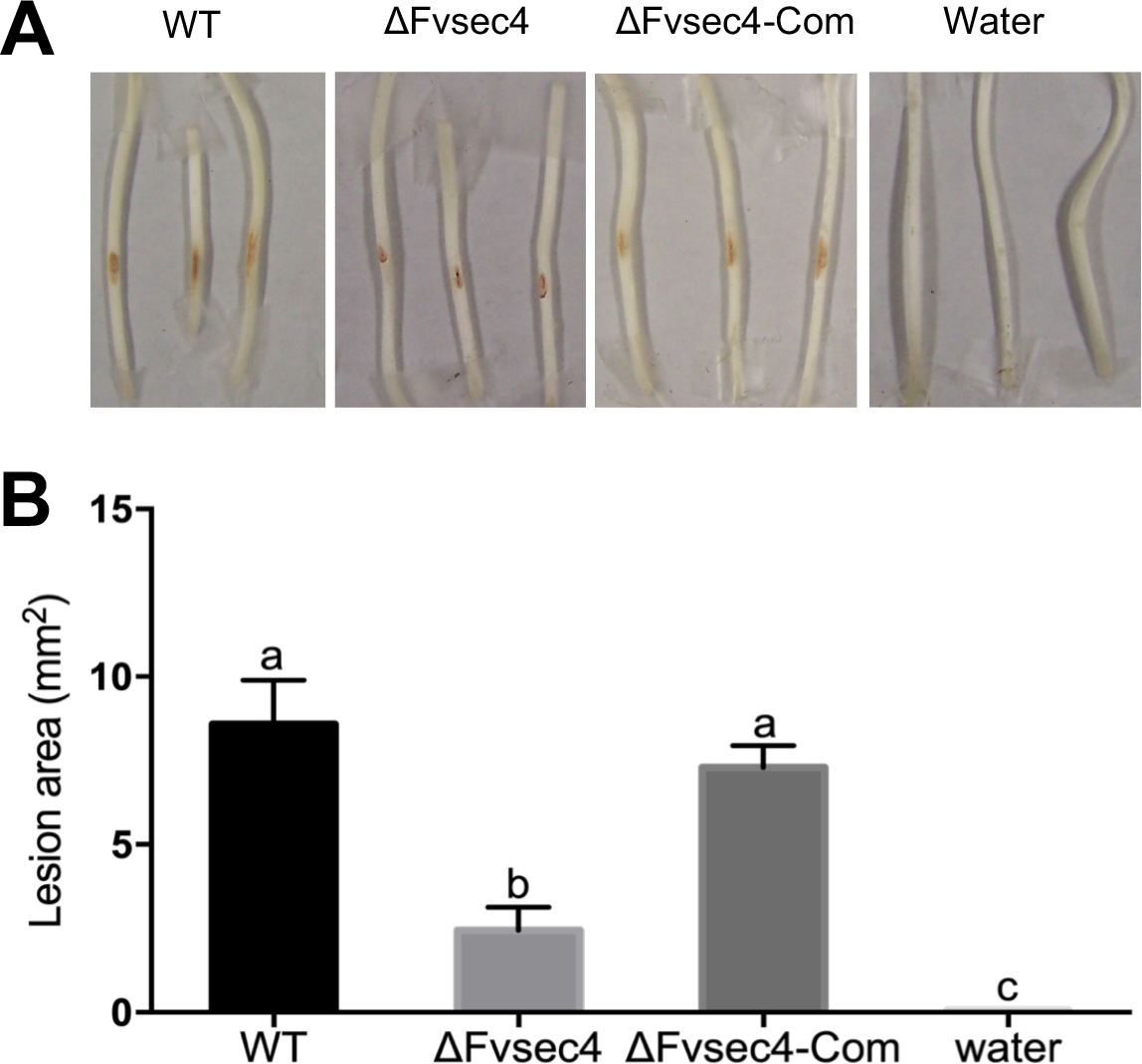
Role of FvSec4 in corn seedling rot severity. (A) We inoculated 10μl of wild-type (WT), ΔFvsec4, and ΔFvsec4-Com spore suspension (10^7^/ml) on one-week old silver queen seedlings. Symptoms were observed after 7 days of incubation. (B) Lesion sizes were quantified using Image J.

### 3.5 FvSec4 is essential for FB1 production

We tested FB1 production in *F. verticillioides* strains on both corn kernels (silver queen hybrid) and in myro liquid medium after one-week incubation. The ΔFvsec4 growth was reduced on corn kernels but not in myro liquid medium (Fig. 6A and S3B). When FB1 production was normalized to fungal growth, the results showed that ΔFvsec4 produces dramatically lower levels of FB1 than the wild-type progenitor in both growth conditions (Fig. 6B and 6C). To further understand how FvSec4 impacts FB1 production at the molecular level, we used qPCR to study the expression of two key FUM cluster genes *FUM1* and *FUM8*. RNA samples were collected from 7 day-post-inoculation myro cultures. Both *FUM1* and *FUM8* expressions were significantly altered with three-fold reduction in ΔFvsec4 strain when compared to the wild-type and ΔFvsec4-Com (Fig. 6D). Additionally, we also tested *SEC5*, *EXO70*, *SYN1* and *LCP1* gene expression in 7 day-post-inoculation myro and 20h YEPD liquid media. Our data showed selected genes associated with exocytosis were not altered but LCP1 transcription level was suppressed in the ΔFvsec4 deletion mutants (Fig. 6D and S3D). Taken together, our results indicate that FvSec4 is positively associated with key *FUM* gene expression and ultimately FB1 production.

**Fig. 6.**
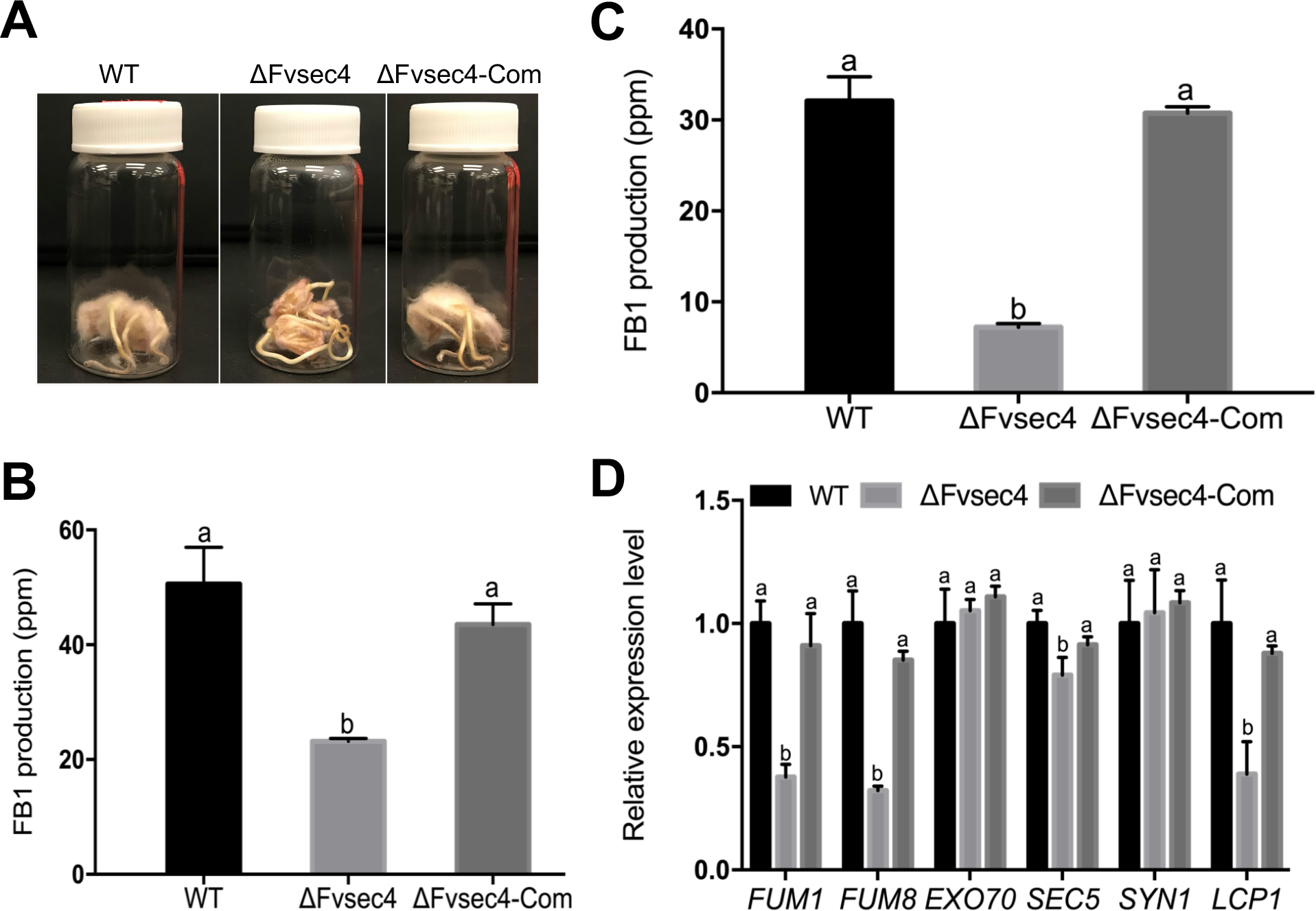
Influence of FvSec4 in FB1 production and key FUM gene transcription. (A) Surface sterilized silver queen corn seeds were inoculated with wild-type (WT), ΔFvsec4, ΔFvsec4-Com and incubated for 7 days. Sterile water was used as a negative control. (B) FB1 and ergosterol were quantified by HPLC. Ergosterol level in each sample was used to normalize FB1 levels, thus resulting in relative FB1 production in corn seeds. (C) Myro liquid medium was inoculated with WT, ΔFvsec4, and ΔFvsec4-Com for 7 days at room temperature with agitation. FB1 levels were analyzed by HPLC. (D) Transcriptional analyses of key FUM genes, exocyst-related genes, and FvLcp1 in WT, ΔFvsec4, ΔFvsec4-Com after 7-day incubation in the myro liquid medium. Transcripts were normalized against WT gene expression. Three biological replicates were performed independently. Error bars in this study all represent the standard deviations for three replicates. Lowercase letters in this study on the bar top suggest significant differences among various strains.

### 3.6 FvSec4 plays an important role in response to various stressors and extracellular enzymes secretion

To investigate whether FvSec4 is involved in response to environment stress agents, we tested vegetative growth of strains on minimal media amended with SDS (cell wall stress), Congo red (cell wall stress) and H_2_O_2_ (oxidative stress) (Fig. 7A). The mutant growth rate was significantly inhibited by these stress agents when compared to the wild-type strain, especially under the oxidative stress with H_2_O_2_. (Fig. 7B). This outcome suggests that FvSec4 plays a role in response to stress related to cell wall integrity and tolerance to oxidative stress.

**Fig. 7.**
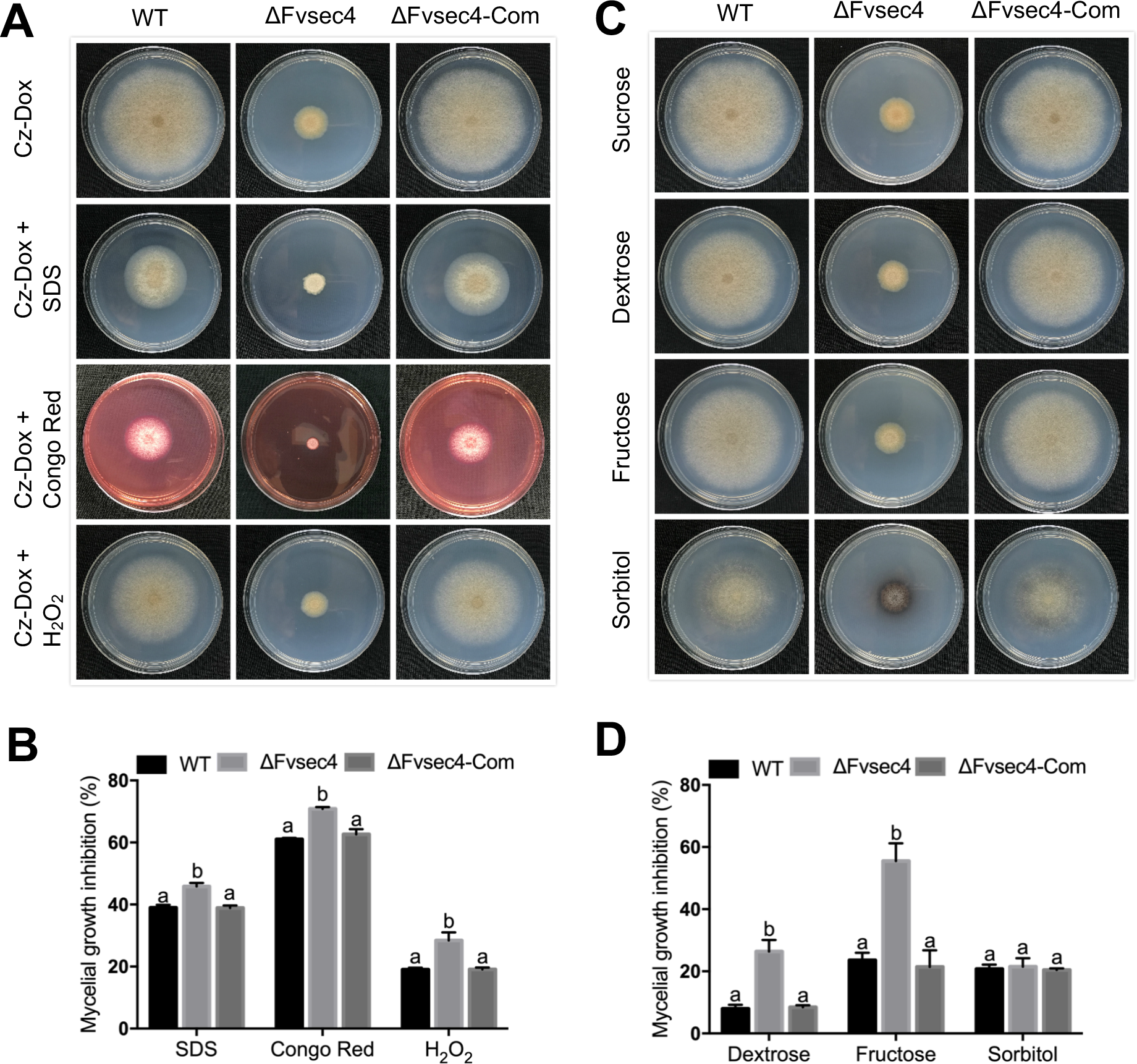
Susceptibility against cell wall stress agents and deficiency in carbon utilization in ΔFvsec4 mutant. (A) Strains were grown on Czapek-Dox agar amended with SDS, congo red and H_2_O_2_ for 8 days at room temperature. (B) Growth diameter of ΔFvsec4, ΔFvsec4-Com were subjected to statistical analyses. The growth inhibition rate (%) was measured by (sucrose growth diameter - designated stress growth) / sucrose growth diameter x 100. (C) Strains were grown on modified Czapek-Dox with dextrose, fructose or sorbitol as the sole carbon source for 8 days at room temperature. Czapek-Dox agar plates with sucrose was used as a control. (D) Inhibition rate of strains grown on the media containing different carbon sources. Three replicates were used in this assay. The growth inhibition rate (%) was measured by (diameter of growth on sucrose - designated carbon growth) / diameter of growth on sucrose x 100.

To determine if FvSec4 is important for utilization of different carbon nutrients, we cultivated wild-type, ΔFvsec4, ΔFvsec4-Com strains on Czapek-Dox agar medium containing different carbon sources, *i.e.* sucrose, dextrose, fructose and sorbitol (Fig. 7C). We learned that the mycelial growth of ΔFvsec4 mutant exhibits significant restriction when grown on Czapek-Dox agar containing dextrose or fructose but not sorbitol, when compared to the growth observed with sucrose as the sole carbon source (Fig. 7D). Further studies are needed to determine if this deficiency is due to carbon nutrient import into fungal cell or secretion defect in extracellular catabolic enzymes.

## Discussion

Exocytosis plays important roles in diverse functions such as cell polarization, growth, morphology, and migration (He and Guo 2009). Exocytosis is responsible for the secretion of cellular substances to the extracellular space. When pathogenic fungi colonize the living plants, these organisms employ various strategies to adapt the host environment, namely by activating signaling pathways associated with producing effectors, secondary metabolites and enzymes (van der Does and Rep 2017). Previous studies indicate the Sec4 protein is a key regulator of multi-subunit exocyst complex function (Guo, et al. 1999). Unlike in *S. cerevisiae* and *Candida albicans*, Sec4 protein does not appear to be essential for viability but its function is critical for other physiological functions in filamentous fungi (Salminen and Novick 1987; Mao, et al. 1999). The deletion of FvSec4, a highly conserved Rab GTPase protein, showed that this protein is essential for the hyphal branching and growth, conidiation, stress responses and extracellular enzymes secretion in *F. verticillioides*. FvSec4 was also indispensable for the virulence and mycotoxin production.

Similar to ΔMosec4 in *M. oryzae*, the virulence in ΔFvsec4 strain was significantly reduced in our seedling rot assay when compared with the wild-type progenitor. The reduced virulence in ΔMosec4 was partially due to appressoria abnormalities, particularly with lower turgor pressure which is crucial for host penetration. Misshapen invasive hyphae and mislocalization of the cytoplasmic effector in ΔMosec4 mutant could have also negatively impacted virulence. Infection structure such as appressoria are not recognized in *F. verticillioides.* But it is also noteworthy that a mutation in a Sec4 homolog in wheat scab pathogen *F. graminearum*, a non-appressorium producing ascomycete, also led to a virulence defect (Zheng, et al. 2015). Furthermore, the deletion of Sec4 homolog BcSas1 in *Botrytis cinerea* also showed reduced virulence, and this outcome could perhaps be explained by reduced growth rate and inadequate secretion of cell wall degrading enzymes (Zhang, et al. 2014). While we cannot exclude slower vegetative growth as one of the factors for reduced virulence, we can also propose that FvSec4 is involved in regulating the expression of *F. verticillioides* secreted proteins. Consistent with this idea, we learned that the expression of *FvLCP1* gene was significantly decreased in ΔFvsec4. In our previous study, we characterized FvLcp1 as a secreted protein that is involved host defense suppression and FB1 biosynthesis (Zhang, et al. 2017).

Another possible reason for attenuated pathogenicity is due to the mutant exhibiting deficiencies in responding to various exogenously applied stress agents. In our study, ΔFvsec4 showed increased sensitivity to H_2_O_2_. It is well studied that reactive oxygen species (ROS) are accumulated in plant hosts as a response to biotic and abiotic stress, and *Fusarium* species are directly causing biotic stress on corn (Lehmann, et al. 2015). However, there are contradicting studies that suggest ROS sensitivity may not be a key factor in fungal virulence. For instance, *B. cinerea* BcSas1 deletion mutants showed less sensitivity when compared to the wild-type (Zhang, et al. 2014). In addition to ROS response, ΔFvsec4 exhibited growth deficiencies when utilizing different carbon sources such as dextrose and fructose when compared to sucrose. This result raises a question whether FvSec4 is involved in secretion of enzymes important for specific carbon nutrient utilization. The hypersensitivity to H_2_O_2_ and the impairment in carbon nutrient utilization are consistent with the attenuated virulence in ΔFvsec4. Published studies show that Sec4 Rab GTPase are important for stress response and nutrient utilization (Zhang, et al. 2014; Zheng, et al. 2016). But, whether these two physiological processes are genetically linked needs further investigation.

Secondary metabolites are not required for conventional growth in *Fusarium* species but may offer advantages in certain circumstances (Ma, et al. 2013). However, it is clear that mycotoxins produced by fungi have adverse effects on human health and food safety (Wu, et al. 2014). There is an earlier study by Zheng et al (2015) describing how Rab GTPase FgRab8 and exocytosis are positively associated with *F. graminearum* mycotoxin DON production (Zheng, et al. 2015). While DON has been recognized as a virulence factor in *F. graminearum*, FB1 is not a critical factor for plant pathogenesis in *F. verticillioides* (Proctor, et al. 1995; Desjardins, et al. 2002). In this study, ΔFvsec4 showed a significantly lower levels of FB1 production when compared to the wild-type strain, implying that this protein is critical for regulating mycotoxin biosynthesis. To further understand how FvSec4 is impacting FB1 production, we tested the expression of key FUM genes *FUM1* and *FUM8*. Our qPCR result showed that transcriptional expression of these two FUM genes were significantly suppressed in the mutant. It is reasonable to hypothesize that this Rab GTPase indirectly regulate transcriptional activities of FUM cluster through other transcription factors.

Sec4 protein is known to control exocyst assembly and involved in secretion of vesicles. We confirmed that FvSec4 is localized to the hyphal tip area which is consistent with the exocytosis function. To investigate the role of FvSec4 in regulating the exocyst complex and other downstream components, we identified three Sec4 downstream genes, *EXO70*, *SEC5* and *SYN1*, to study their gene expression levels. We found that ΔFvsec4 mutation is not crucial for exocyst-related gene expression except for *SEC5*, which showed 22% less expression when compared to wild-type progenitor after a 7-day incubation in myro medium. This outcome led us to conclude that Sec4 is not directly involved in the transcriptional regulation of its downstream genes associated with exocytosis.

Spitzenkörper is a subcellular structure found at the fungal hyphal tip that is associated with polar growth (Riquelme and Sanchez-Leon 2014). A previous report also indicated that Spitzenkörper acts as a Vesicle Supply Center (VSC) where vesicles accumulate before being released to extracellular space (Bartnicki-Garcia, et al. 1989). We stained mycelia with FM4-64, and the result showed that our wild-type strain harbors a recognizable Spitzenkörper at the hyphal tip. However, ΔFvsec4 exhibited accumulated fluorescence in both apical and subapical areas while the Spitzenkörper was absent after the same staining treatment. When we consider the important role of Spitzenkörper in delivering cell wall components to the sites of cell wall synthesis, perhaps this abnormal Spitzenkörper structure and distribution are impacting the response to SDS and congo red cell wall stress agents in ΔFvsec4 (Riquelme 2013). The lack of Spitzenkörper is possibly due to ΔFvsec4 showing a defect in maintaining the balance between exocytosis and endocytosis. We can further hypothesize that in the mutant insufficient number of vesicles are delivered to the hyphal tip, and perhaps this leads to a significantly slower vegetative growth and hyper-branching.

## Acknowledgements

We thank Dr. Brian Shaw, Ms. Blake Commer and Mr. Joe Vasselli (Department of Plant Pathology and Microbiology, Texas A&M University) for help and discussion in microscopy. This research was supported in part by the Agriculture and Food Research Initiative Competitive Grants Program Grant (2013-68004-20359) from the USDA National Institute of Food and Agriculture. The authors declare no conflict of interest.

---

**Fig.S1.**
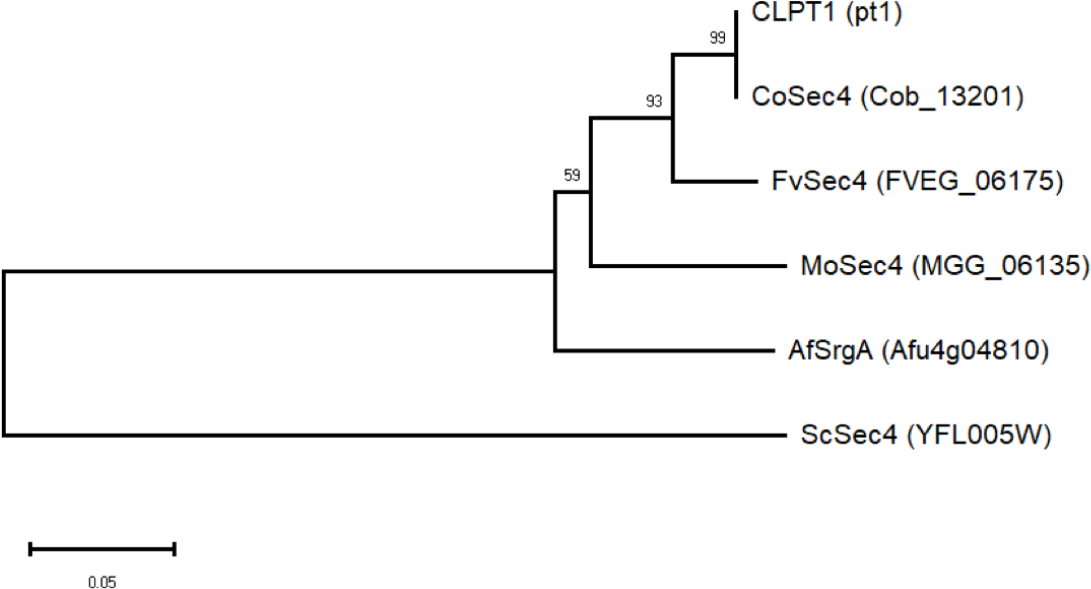
(A) Phylogenetic analysis for FvSec4 and other Sec4 orthologs. Multiple sequences were aligned by ClusterX2. A neighbor-joining tree was constructed by MEGAX. The number at nodes indicates the percentage of replicate trees clustered together in 1000 bootstrap replications.

**Fig.S2.**
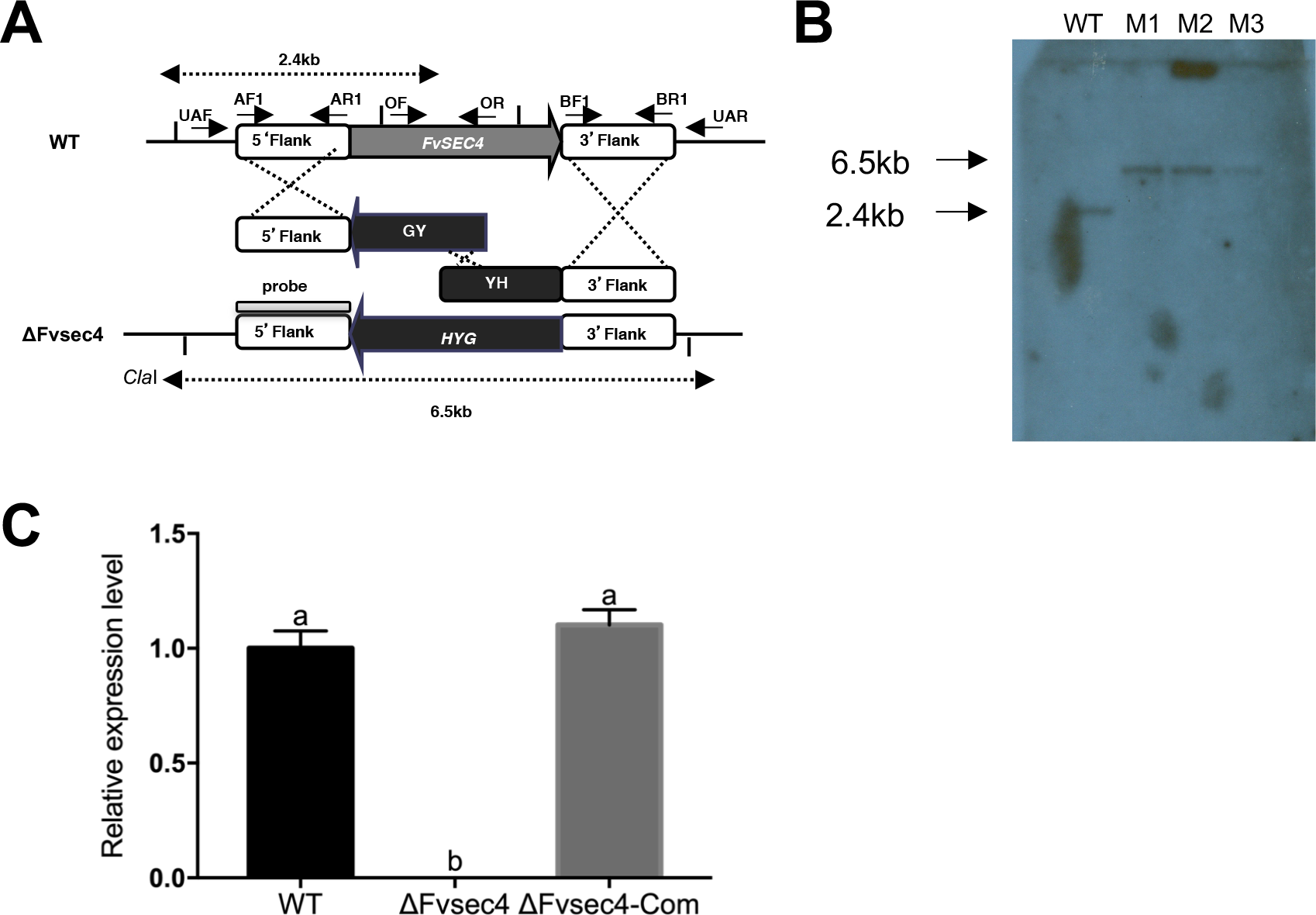
(A) Homologous gene recombination approach used to construct ΔFvsec4 mutant in *F. verticillioides*. (B) Southern blot analyses of WT and three knockout candidates (M1, M2, M3). M1 was used in this study as ΔFvsec4. We used the 5’-flanking regions as a probe for Southern blot. (C) WT, ΔFvsec4 and ΔFvsec4-Com total RNAs were isolated after 7 days in myro liquid medium and reversed to cDNA. qPCR was employed to verify the ΔFvsec4 and ΔFvsec4-Com using a *FvSEC4* gene-specific primer set while WT was as a control. No detectable FvSec4 transcript in the ΔFvsec4 mutant by qPCR analysis and ΔFvsec4-Com fully recover the *FvSEC4* expression.

**Fig.S3.**
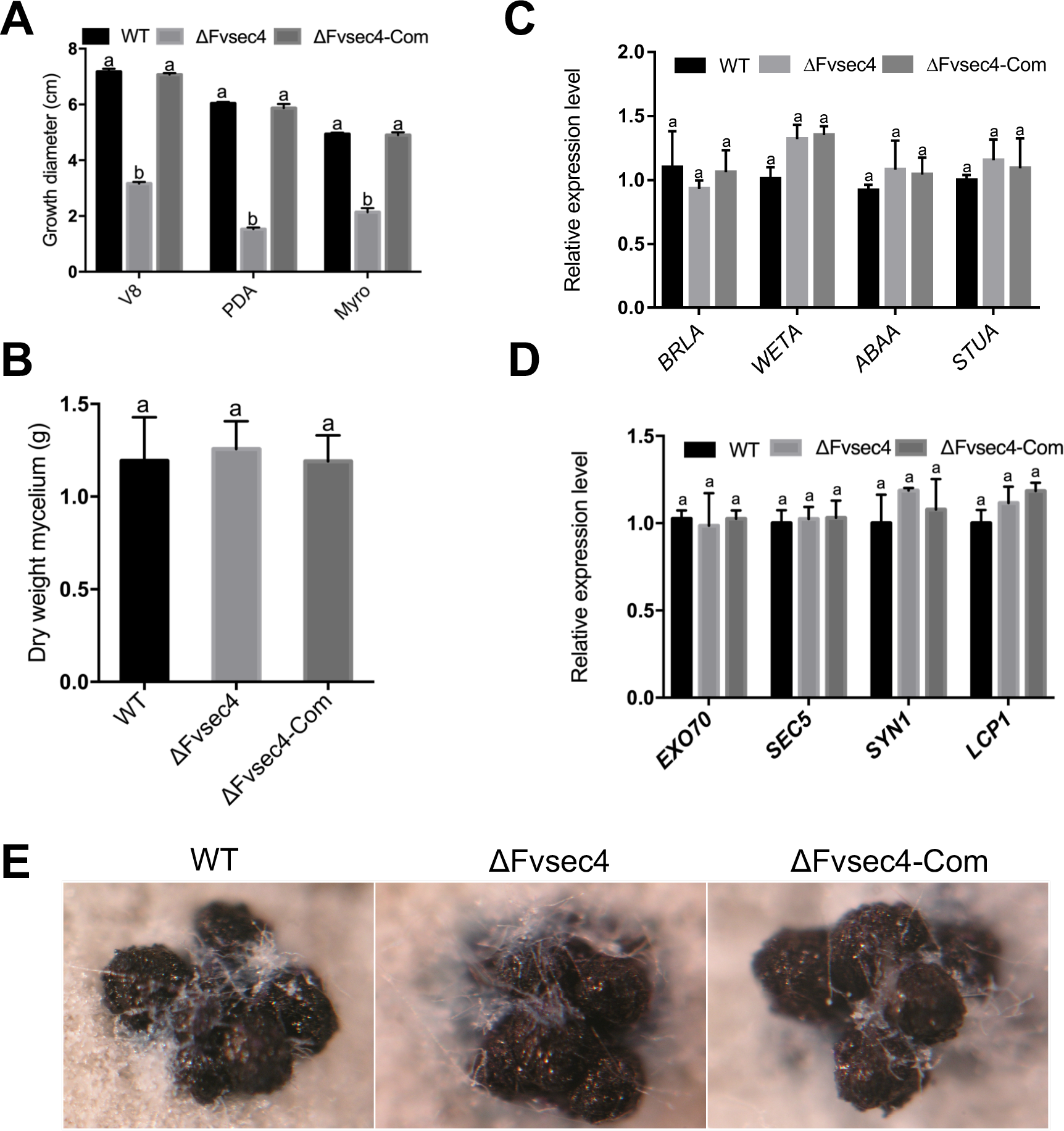
Involvement of FvSec4 the vegetative growth but not perithecia in sexual mating. (A) Mycelia growth diameter on V8, 0.2xPDA and myro solid medium were assayed after 8 days at room temperature. (B) Same weight of mycelia from YEPD was transferred into myro liquid medium for constant shaking. Samples were collected after 7 days incubation in 100ml myro liquid medium. (C) qPCR study of *BRLA*, *WETA*, *ABAA* and *STUA* after 20 h incubation in YEPD liquid medium (D) 20 h samples in YEPD medium was studied the expression of exocyst-related and FvLcp1 gene expression. (E) WT, ΔFvsec4 and ΔFvsec4-Com were crossed to M3120, an opposite mating type to assay fertility.

**Table S1.**
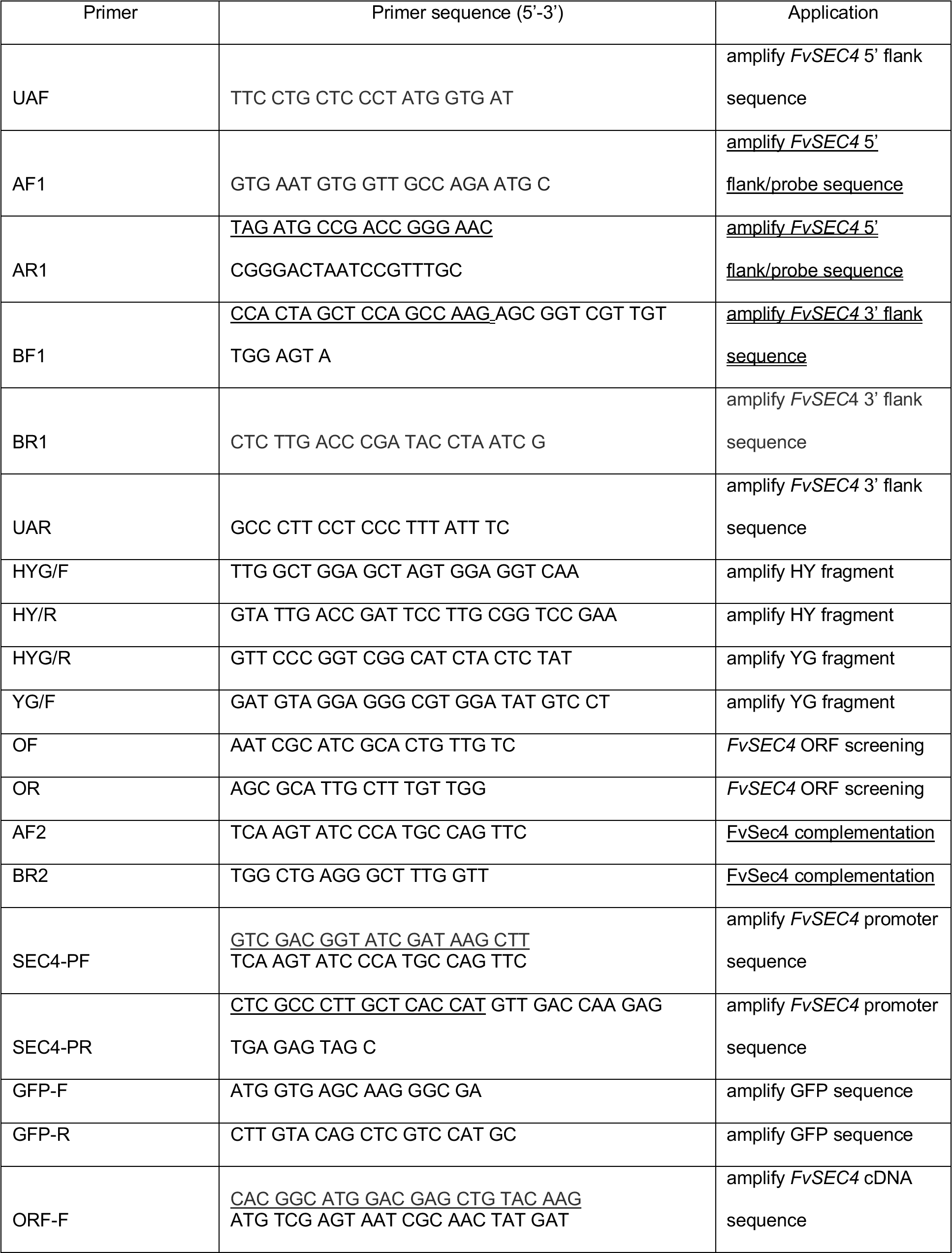
Primers used in this study

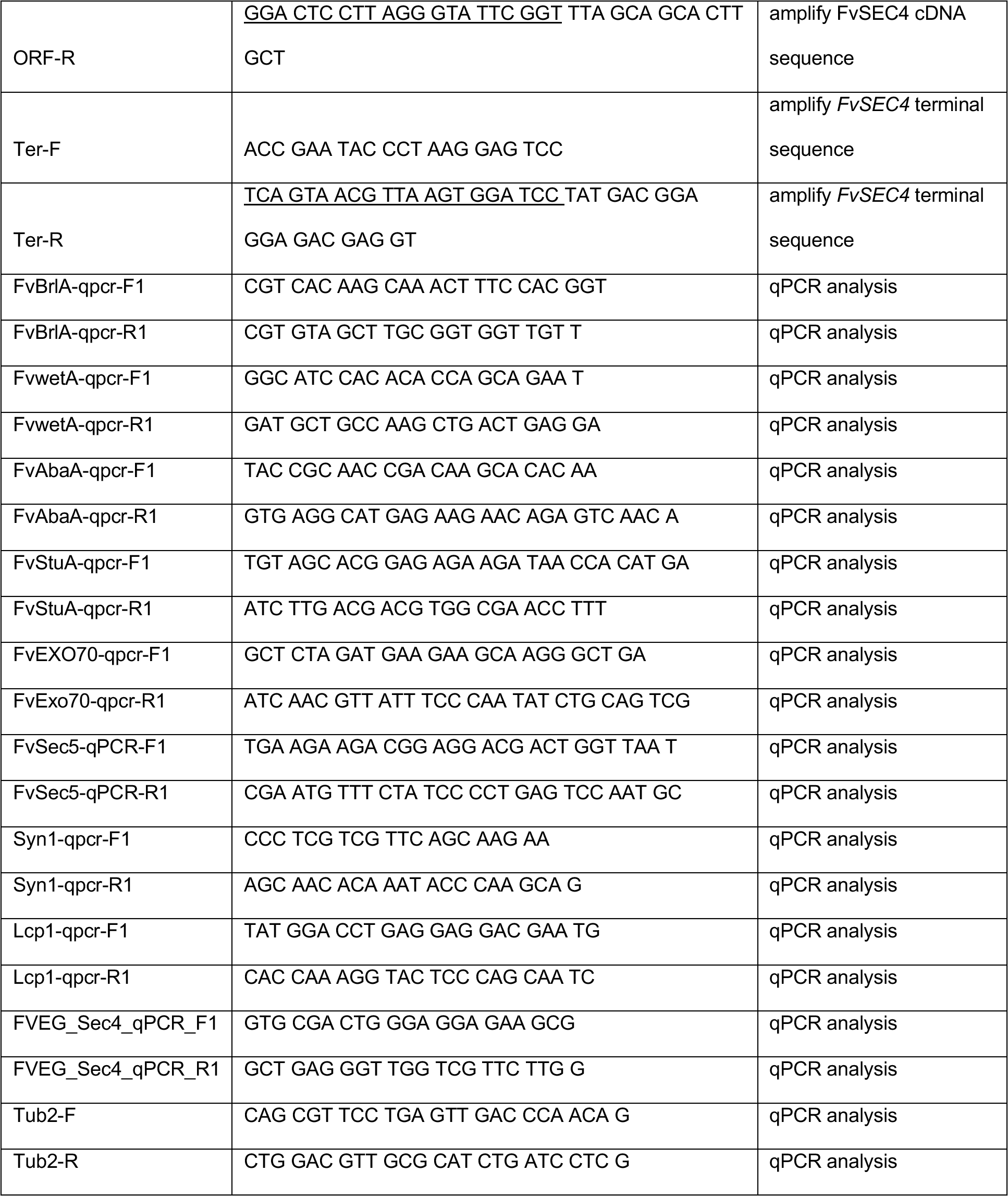

